# Finite-sample genome-wide regression p-values (GWRPV) with a non-normally distributed phenotype

**DOI:** 10.1101/204727

**Authors:** Gregory Connor, Michael O’Neill

## Abstract

This paper derives the exact finite-sample p-value for univariate regression of a quantitative phenotype on individual genome markers, relying on a mixture distribution for the dependent variable. The p-value estimator conventionally used in existing genome-wide association study (GWAS) regressions assumes a normally-distributed dependent variable, or relies on a central limit theorem based approximation. The central limit theorem approximation is unreliable for GWAS regression p-values, and measured phenotypes often have markedly non-normal distributions. A normal mixture distribution better fits observed phenotypic variables, and we provide exact small-sample p-values for univariate GWAS regressions under this flexible distributional assumption. We illustrate the adjustment using a years-of-education phenotypic variable.

## 1 Introduction

This paper provides an alternative estimator of the coefficient p-values in genome-wide univariate regressions of a phenotypic variable on a single-nucleotide polymorphism (SNP). The formula is easy to apply, and can provide substantially more accurate p-values if the phenotypic variable under consideration is non-normally distributed and the number of observations is not extremely large. For a normally distributed phenotypic variable, or with an extremely large sample, the adjustment is not necessary. The magnitude of the p-value adjustment depends upon the size of the sample, the non-normal features of the phenotypic variable including skewness and kurtosis, and the minor allele frequency of the SNP.

Regression based Genome-Wide Association Studies (GWAS) are a key exploratory tool in genetic research on the heritability mechanisms of phenotypic traits, with the goal of identifying individual SNPs associated with a trait. GWAS involves a million or more individual regressions (one per SNP), in the search for SNPs with a significant relationship to an observed phenotypic variable. To account for the multiple comparisons problem, analysts use Bonferonni-corrected p-values, so that an adjusted 5% p-value with one million independent regressions requires an uncorrected univariate regression coefficient p-value (for a two-sided test) of 0.025*x*10^−6^.

In the estimation of Bonferonni-corrected p-values, analysts rely on the assumption that the estimated regression coefficient is normally distributed. This holds exactly if the phenotypic variable has a normal distribution, and approximately (for sufficiently large samples) if the phenotypic variable has any reasonably well-behaved distribution, by the central limit theorem. The quality of the central limit theorem based approximation depends upon the size of the sample, the distributional characteristics both of the observed phenotypic variable and the SNP, and (crucially in this application) on the magnitude of the p-value.

The central limit theorem guarantees uniform convergence of the true cumulative distribution to the normal distribution (see White (1984) for a review). An approximate p-value in the region of 0.025, accurate to within ±0.0001, can be entirely adequate; an approximate p-value in the region of 0.025*x*10^−6^ = 0.000000025 which is similarly accurate to within ±0.0001 is effectively worthless. GWAS involve very large-number multiple tests and therefore extremely low p-value thresholds, with the conventional critical value set at 0.025*x*10^−6^. Invocation of the central limit theorem is problematic in this context.

In this paper, we develop an alternative approach based on fitting a Bernoulli-normal mixture distribution to the phenotypic variable. As we show, since a genome-wide regression has a trinomial explanatory variable (the three states of the SNP) and the Bernoulli-normal mixture is a combination of a binomial and a normal, the resulting regression coefficient p-value is a multinomial-based linear combination of independent normals, with a closed-form expression in terms of the standard normal distribution. Empirically, the p-value adjustment can be quite large, and can increase or decrease estimated p-values relative to the conventional approach. We illustrate the approach by comparing conventional and adjusted p-value for years-of-education, a common phenotypic variable which has a notably non-normal distribution.

Assuming that the mixture distribution correctly describes the phenotypic variable, our finite-sample adjusted p-value eliminates the reliance on the large-sample-dependent central limit theorem for significance tests in GWAS. This allows for the use of datasets with modest sample sizes and for including SNPs with very low minor allele frequencies. We provide an implementation of the method in the R programming language, as a standard R package from CRAN, along with user documentation.

## 2 Exact finite-sample p-values for GWAS univariate regression under a Bernoulli-normal mixture distribution

This section presents the new methodological result. We derive the exact p-values of a univariate GWAS regression under an assumed Bernoulli-normal mixture distribution. This is a reasonably straightforward exercise, combining the Bernoulli-normal mixture for the dependent variable with the three-valued explanatory variable of the regression, and then rearranging, manipulating, and simplifying the expressions.

### 2.1 The GWAS regression framework

The analyst has observations on *i* = 1, *n* individuals with the data consisting of a phenotypic variable (such as income, years of education, life satisfaction rating, etc.) and a very long (we assume 10^6^ for notational simplicity) string of genetic markers. We assume that the phenotype variable *y* has known mean and variance; to simplify notation (without loss of generality) we assume that *y* is standardized and so has zero mean and unit variance. We assume that the phenotype variable *y* has been pre-whitened with respect to any other confounding explanatory variables by a preliminary regression step, so that the GWAS regressions are univariate. The genetic markers are single nucleotide polymorphisms which have three potential states: major allele homozygote, minor allele homozygote, and heterozygote. Let *x*_*ij*_ be the trinomial explanatory variable, set equal to 0 if individual i is a major allele homozygote for the *j*^*th*^ genetic marker, 1 if he/she is a heterozygote, and 2 if he/she is a minor allele homozygote.

The research objective is to identify individual SNPs that have an influence on the phenotypic variable. This is done through a set of 10^6^ univariate regressions, each using one genetic marker. As null hypothesis, we assume that none of the individual SNPs have an influence on the phenotype, so that:

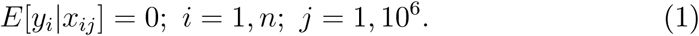

This null hypothesis is tested separately for each *j*. Let *m*_*x*_ and 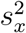 denote the sample average and mean-square deviation of the explanatory variable *x*_*j*_. For each individual *j* we test the statistical significance of the ordinary least squares regression coefficient:

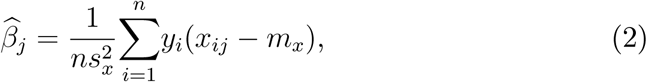

where it follows from (1) that 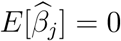 for all *j*.

In performing the multiple tests 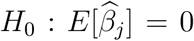 with *j* = 1, .., 10^6^ each tested separately, it is crucial to apply a Bonferonni correction to the individual test critical values. With 10^6^ independent tests, and choosing a 95% confidence level, the two-tailed critical values for Bonferonni-corrected multiple-test significance of each coefficient uses a cumulative probability of 0.025 × 10^−6^ for testing a negative estimated coefficient and 1 − .025 × 10^−6^ for a positive estimated coefficient.

### 2.2 Fitting a Bernoulli-normal mixture distribution to a phenotypic variable

As we will demonstrate later, the central limit theorem does not always provide a reliable approximation for the extremely small p-values under examination in the large-number multiple test environment of GWAS. We need an alternative estimator of regression p-values to apply in the case of a non-normally distributed phenotypic variable. We need a reasonable distributional assumption on *y* that fits the phenotypic variable and allows for the feasible computation of exact p-values that do not rely on the central limit theorem approximation. In this section we propose a Bernoulli-normal mixture distribution.

The Bernoulli-normal mixture distribution is a flexible family of distributions with good fit in many applications, and convenient analytical properties in our model.

Let 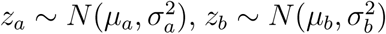, and λ a Bernoulli distributed random variable with λ = 1 with probability *p*; all three random variables assumed independent. The mixed normal *y* is the random variable:

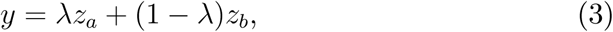

which has five parameters: *µ*_*a*_, *σ*_*a*_, *µ*_*b*_, *σ*_*b*_, *p*. The first two moments are:

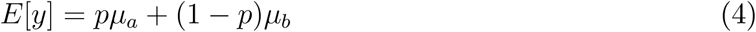

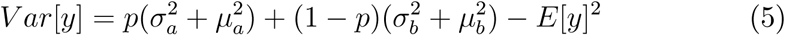

For notational simplicity, consider the case in which *E*[*y*] = 0 and *Var*[*y*] = 1, then the third and fourth moments are:

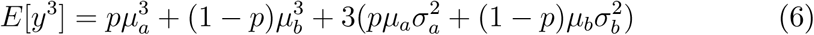

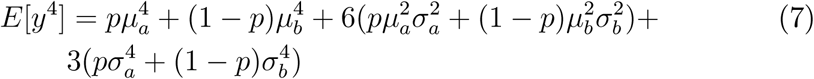

The distribution can be fitted via EM-maximum likelihood; see McLachlan and Peel (2000) for an overview of mixture distributions and estimation methods.

## 3 The GWAS univariate regression coefficient as a linear combination of independent normals

We now use the assumption that *y* has a Bernoulli-normal mixture distribution to derive the exact finite-sample p-values of 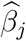. Since we now look at one particular *j* only, we simplify notation and drop the *j* subscript. To implement our technique, the analyst counts the number of major allele observations, heterozygote observations and minor allele observations in each regression sample. Let {*n*_0_, *n*_1_, *n*_2_} denote these three integer values, with *n*_0_ + *n*_1_ + *n*_2_ = *n*. The sample average and mean-square deviation of the explanatory variable have simple forms, since x_i_ only takes the three values 0, 1, 2:

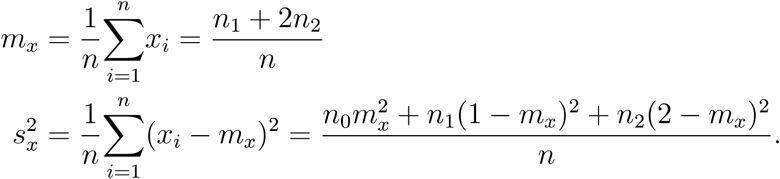

The cumulative distribution of an estimate 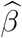 is the probability under the null hypothesis of a random realization 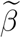 having that value or less:

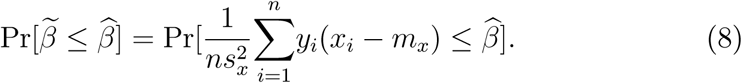

For notational convenience, we order the observations index so that the major allele observations are listed first, then the heterozygote observations, and then the minor allele observations.

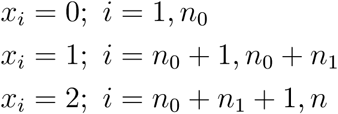

Writing out the coefficient formula (8) using the observed values *n*_0_, *n*_1_, *n*_2_:

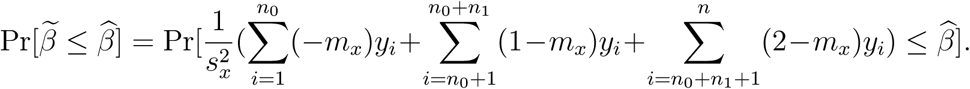

Under our distributional assumption on *y*, each of the three integers *n*_0_, *n*_1_ and *n*_2_ in turn decomposes into two (unobserved) integers: the number of realizations of the dependent variable 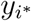 with conditional mean and standard deviation either {*µ*_*a*_, *σ*_*a*_} or {*µ*_*b*_, *σ*_*b*_}. We denote these integer realizations with a double subscript, {*n*_0*a*_, *n*_1*a*_, *n*_2*a*_}_*h*_,and {*n*_0*b*_, *n*_1*b*,_ *n*_2*b*_}_*h*_, *h* = 1, *m* where the first subscript refers to the minor allele frequency, 0, 1, or 2, and the second subscript refers to the (unobserved) number of realizations of {*µ*_*a*_, *σ*_*a*_} or {*µ*_*b*_, *σ*_*b*_}. Since the second set {*n*_0*b*_, *n*_1*b*,_ *n*_2*b*_} consists of the remainders *n*_0*b*_ = *n*_0_ − *n*_0*a*_, *n*_1*b*_ = *n*_1_ − *n*_1*a*_ and *n*_2*b*_ = *n*_2_ − *n*_2*a*_, it is sufficient to index with {*n*_0*a*_, *n*_1*a*_, *n*_2*a*_}.

Each of the integers *n*_0*a*_, *n*_1*a*,_ *n*_2*a*_ has an independent binomial distribution, since each depends upon the number of *λ* = 1 outcomes of the Bernoulli random variable *λ* within one of the three allele categories. It is easy to see that *m* = (*n*_0_ + 1)(*n*_1_ + 1)(*n*_2_ + 1). The probabilities of each of the potential outcomes {*n*_0*a*_, *n*_1*a*,_ *n*_2*a*_}_*h*_, *h* = 1, *m* can be found from the binomial formula (for three independent binomials):

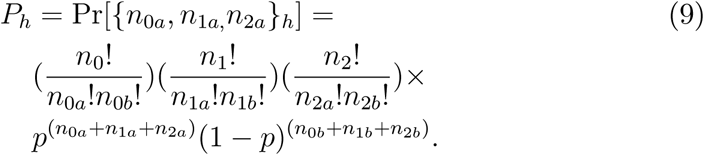

The cumulative probability of 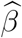 is the sum of the conditional cumulative probability given each of the potential outcomes *h* = 1, *m* times the probability of each outcome:

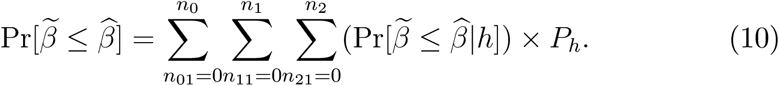

The last step is to calculate

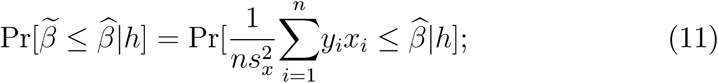

which is the sum of n independent normals, consisting of *n*_0*a*_ draws with mean −*m*_*x*_*µ*_*a*_ and standard deviation of *m*_*x*_*σ*_*a*_, plus *n*_0*b*_ draws with mean −*m*_*x*_*µ*_*b*_ and standard deviation of *m*_*x*_*σ*_*b*_, plus *n*_1*a*_ draws with mean (1 − *m*_*x*_)*µ*_*a*_ and standard deviation of (1 − *m*_*x*_)*σ*_*a*_, plus *n*_1*b*_ draws with mean (1 − *m*_*x*_)*µ*_*b*_ and standard deviation of (1 − *m*_*x*_)*σ*_*b*_, plus *n*_2*a*_ draws with mean (2 − *m*_*x*_)*µ*_*a*_ and standard deviation of (2 − *m*_*x*_)*σ*_*a*_, plus *n*_2*b*_ draws with mean (2 − *m*_*x*_)*µ*_*b*_ and standard deviation of (2−*m*_*x*_)*σ*_*b*_. A sum of independent normals has a normal distribution with mean equal to the sum of the means and variance equal to the sum of the variances. Applying this to (11):

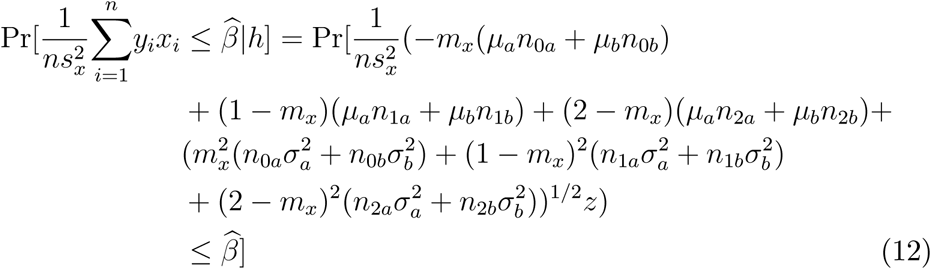

where *z* denotes a standard normal random variable. The right-hand side of equation (12) is simply the p-value of a normally distributed random variable. Therefore, the p-value of the regression coefficient (10) is a binomial-weighted linear combination of conventional normal distribution p-values. There is no need to invoke the (sometimes unreliable) central limit theorem approximation to find the regression p-value. Given the trinomial set of values for the independent variable and a Bernoulli-normal mixture distribution for the dependent variable, the exact finite-sample p-value of the GWA regression coefficient is a directly computable linear combination of normal distribution p-values; we simply combine (9), (10) and (12).

As a tangential benefit, the model also provides exact formulas for the skewness and kurtosis of the estimated regression coefficient (under the null hypothesis that 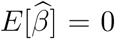. Since *y* is standardized, it also follows from (2) that 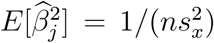. Simply inserting (2) and (3) into the definitions of skewness and kurtosis, and evaluating using the skewness and kurtosis of the mixture distribution, (6) and (7), give closed form expressions that are easy to evaluate, in particular:

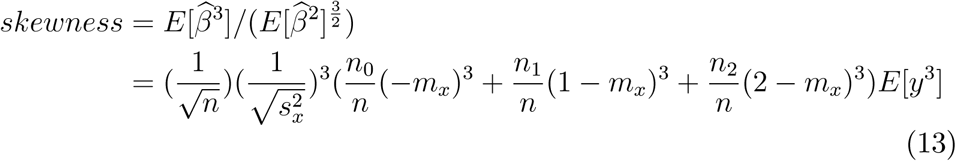

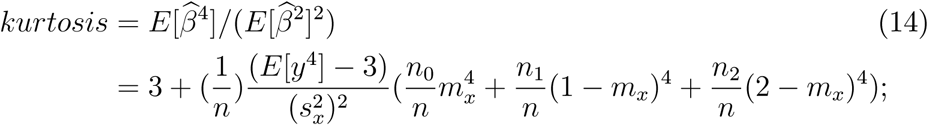

see the Technical Appendix for details. This allows the analyst to have a sense of how close to normality is the finite-sample distribution of the estimated regression coefficient in a particular application.

### 3.1 Computationally efficient implementation of the estimator

The p-value formula (10) requires a sum over the set of outcomes from three independent binomials with *n*_0_, *n*_1_ and *n*_2_ draws, giving a total of *m* = (*n*_0_ + 1)(*n*_1_+1)(*n*_2_+1) terms. For large *n*, the number of terms *m* can be extremely large, but using an efficient computation algorithm, the vast majority of the terms can be dropped from the calculation without any discernible effect on the quality of the estimate. Each of the three random outcomes, *n*_*ja*_, *j* = 0, 1, 2, has an independent binomial distribution with parameter *p* and draw size *n*_*j*_. The three univariate binomial distributions for *n*_*ja*_, *j* = 0, 1, 2 have their peaks at *pn*_*j*_, *j* = 0, 1, 2. The probability of a particular value *n*_*ja*_ diminishes exponentially towards zero as *n*_*ja*_ moves further away from *pn*_*j*_. For large *n*_*j*_, one or both of the "tails" of the univariate probability distribution of *n*_*ja*_ can be deleted, when one or both make approximately zero contribution to the sum (10). See the Technical Appendix for details of the computation algorithm used in the GWRPV estimation code.

## 4 Illustration of the magnitude of the p-value adjustment using years of education

This section examines the magnitude of the adjustment arising from our finite sample p-values compared to using large-sample approximate p-values based on the central limit theorem. We illustrate the adjustment with a commonly used phenotypic variable: years of education, which is a social trait with a strongly non-normal distribution.

Given the parameters of the mixture distribution, our p-value formula (10) is exact; it does not require any simulation. The only inputs needed are the number of major allele homozygote, heterozygote, and minor allele homozygote observations in the regression sample, (*n*_0_, *n*_1_, *n*_2_), the estimated regression coefficient, 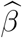, and the five parameters of the mixture distribution, (*p*, *µ*_*a*_, *µ*_*b*_, *σ*_*a*_, *σ*_*b*_).

For the purposes of this comparison, we use five sample sizes, *n* = 500, 1000, 5000, 50000, and 100000. For each sample size we fit *n*_0_, *n*_1_, *n*_2_ from the range of values typically encountered in genome-wide regression tests. Let *MAF* denote the minor allele frequency of the SNP; we chose four representative values: *MAF* = 0.1%, 0.5%, 1%, 5%. To choose the observation numbers *n*_0_, *n*_1,_ *n*_2_ we assume that the SNP is in Hardy-Weinberg equilibrium, which implies that *n*_0_ = *n*(1 − *MAF*)^2^, *n*_1_ = 2*nMAF*(1 − *MAF*) and *n*_2_ = *nMAF*^2^. The numbers of observations *n*_0_, *n*_1_, *n*_2_ must be integers, so for fractional values we stick the "leftover" one or two observation(s) in the heterozygote category.

Note that the relative numbers of explanatory variables across the three categories, *n*_0_, *n*_1_, *n*_2_, can affect the quality of the central limit theorem approximation. For example, with *MAF* = 0.1%, only 0.0001% of SNP observations take the value *x* = 2 (zero observations in most small samples); 0.1998% take the value 1, and for 99.8% of the regression sample, *x* = 0. This unbalanced distribution impacts the speed at which the central limit theorem acts upon the probability distribution of the coefficient estimate, and the asymmetry (right-tail probability versus left-tail probability) of its finite-sample distribution, unless the dependent variable is exactly normal. Our p-value adjustment is particularly valuable for genome-wide regressions with low minor allele frequency since it provides a finite-sample test statistic when the large-sample approximation is unreliable. This will become clear in the tables and figures below.

To calibrate the parameters of the mixture distribution, (*p*, *µ*_*a*_, *µ*_*b*_, *σ*_*a*_, *σ*_*b*_), we run EM-maximum likelihood on the phenotypic variable; see below for details.

### 4.1 Application to a non-normal phenotype: Years-of-education

In this subsection we calibrate the Bernoulli-normal mixture using data on years of education from the U.S. Census Bureau Current Population Survey of Educational Attainment, 2015. See Rietveld, et al. (2013, 2015), Okbay et al. (2016), and references therein for details on the considerable number of GWA regression studies with years-of-education as the phenotypic variable.

Figure 1 shows a frequency distribution of the years-of-education data, along with fitted normal and Bernoulli-normal mixture distributions. See the Technical Appendix for description of the U.S. census data. The mixture distribution picks up the high-peakedness and asymmetry in the data distribution, associated with the 76% frequency of data observations in the range 12 − 16 years, and the secondary clump of observations in the 0 − 6 years range with frequency 3.04%. These features are missed by the fitted normal. The data has skewness of −0.676781 and kurtosis of 5.126954, which both differ significantly from the normal distribution values (zero and three, respectively) with 99% confidence. The Jarque-Bera test rejects normality with 99% confidence.

**Figure 1.**
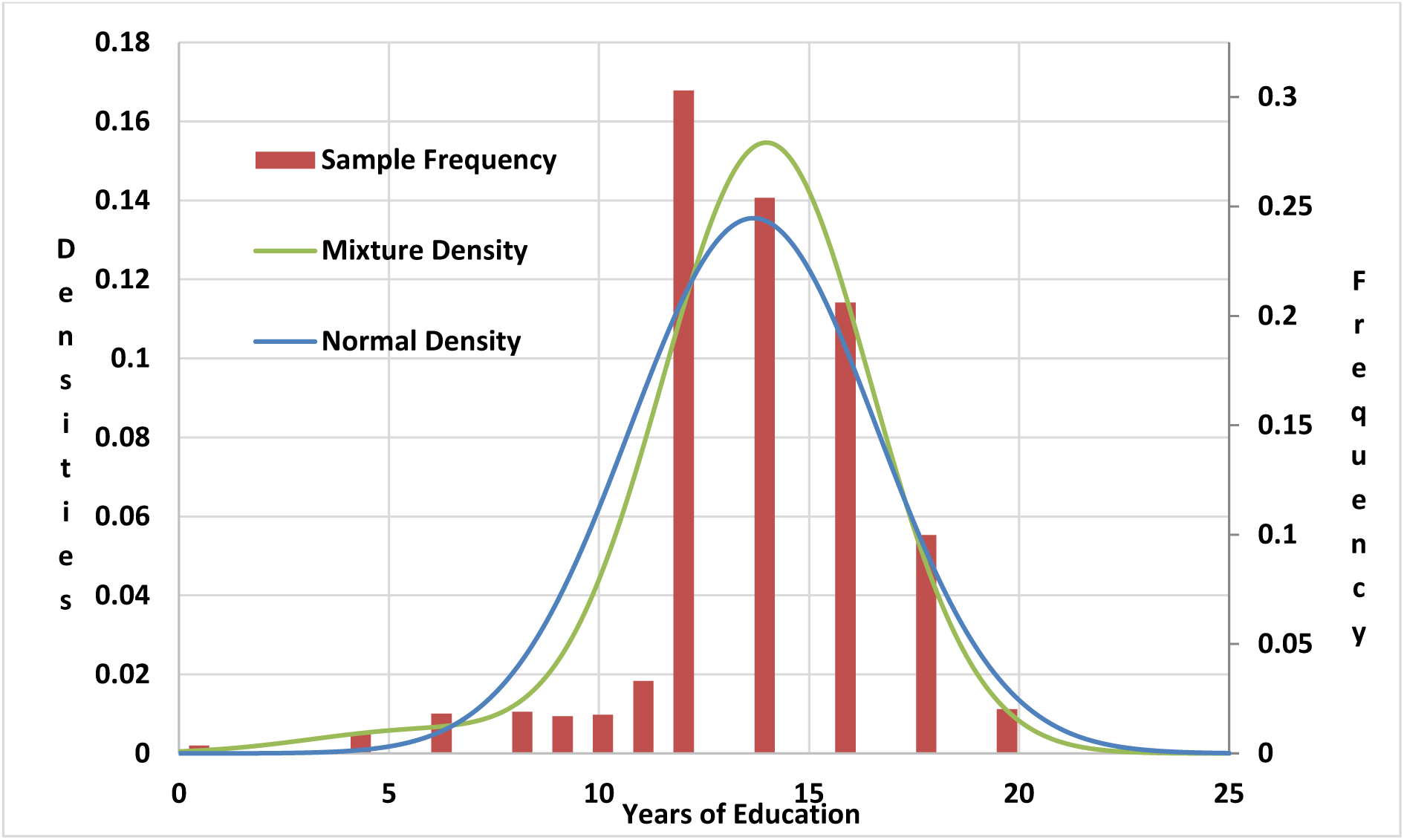
The frequency distribution of years-of-education and fitted normal and mixture distributions

The 81,913 years-of-education data observations are fitted to a Bernoulli-normal mixture distribution using the *normalmixEM* command in the *mixtools* library of programming language *R*; see Benaglia et al. (2009) for details on the estimation routine. The estimated parameter values are *p* = 0.9654, *µ*_*a*_ = 13.872, *µ*_*b*_ = 4.628, *σ*_*a*_ = 2.588, *σ*_*b*_ = 2.518.

For comparative purposes, in the tables below we assume 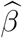 values at the normal-distribution critical values for 95%, and 99% confidence, and for Bonferonni-adjusted multiple-test 95% confidence with 10^6^ independent tests. These 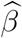 values are easily derived from the t-statistics, using 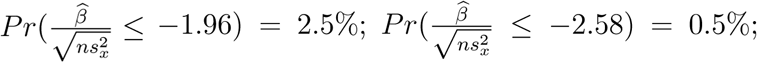 and 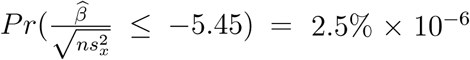; the upper-tail tests are analogous, with a positive sign. We take each of the normal-distribution critical values of 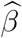 and find their p-values under the mixture distribution, which we can then compare to the normality-based p-values, 2.5%, 0.5%, and 2.5% × 10^−6^.

Table 1 Panel A considers a single-hypothesis, two-sided test with a 95% confidence limit. The table shows exact p-values under the mixture distributions for estimated regression coefficients with normality-based approximate p-values (using the central limit theorem) of 2.5%. The central limit theorem approximation gives quite accurate p-values in almost all cases, even with small sample sizes and low minor allele frequencies. The approximation error from invoking the central limit theorem to estimate p-values is never severe.

**Table 1.**
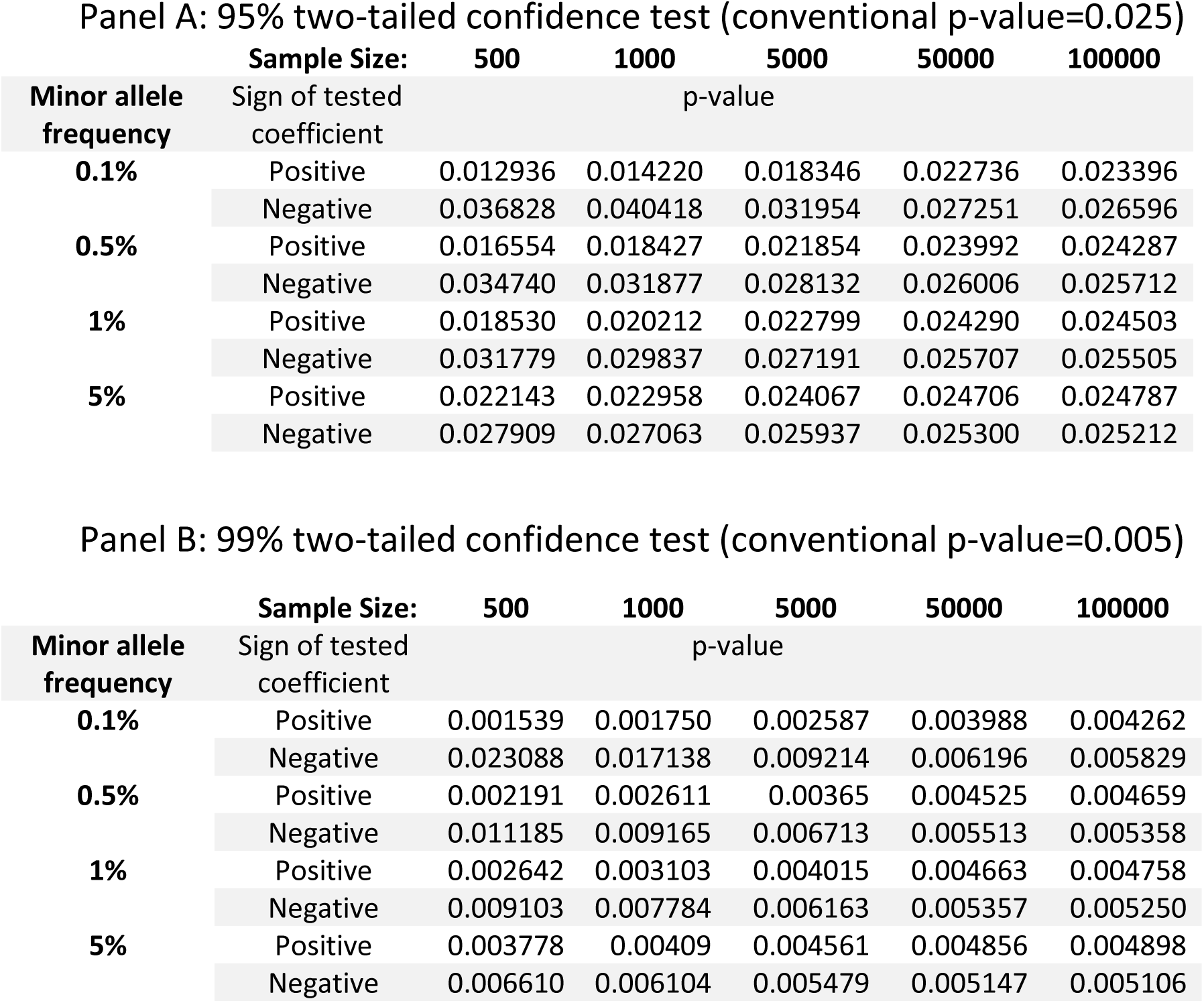
Comparison of adjusted/unadjusted single-test p-values for GWA regression coefficients under a mixture distribution: Years of education

Panel B of the table repeats the exercise for a 99% confidence test, so that the p-value under normality is 0.5%. The central limit theorem approximation continues to work reasonably well, with the exception of smallest sample size (500 observations) with minor allele frequencies of 1% or less.

For 10^6^ multiple test Bonferonni-corrected p-values with 95% confidence, shown in Table 2, the approximation error from relying on the central limit theorem is severe. Convergence of the p-value toward its normality-derived value is much slower, and the small-sample asymmetry in the approximation error is more notable. For small to medium sample sizes, the true p-value for a negative-tail test is very substantially above 2.5% × 10^−6^, the p-value for the positive-tail test is substantially below 2.5% × 10^−6^, and the sum of the two tail probabilities (which should be 5% × 10^−6^) is substantially higher. The central limit theorem approximation only works reasonably accurately for sample sizes of ten thousand or more, and only with relatively high minor allele frequency. In the other cases considered in Table 2, the finite-sample adjustment is critically important.

**Table 2.** Comparison of adjusted/unadjusted 10^6^ multiple-test p-values for GWA regression coefficients under a mixture distribution: Years of education

95% two-tailed confidence test (conventional $p\text{-value} \times 10^6 = 0.025$ )
| Sample Size: |  | 500 | 1000 | 5000 | 50000 | 100000 |
| --- | --- | --- | --- | --- | --- | --- |
| Minor allele frequency | Sign of tested coefficient | $p\text{-value} \times 10^6$ | | | | |
| 0.1% | Positive | 0.000086 | 0.000115 | 0.000349 | 0.003322 | 0.005567 |
|  | Negative | 28.85998 | 56.65438 | 4.016193 | 0.199445 | 0.112986 |
| 0.5% | Positive | 0.000213 | 0.000365 | 0.001747 | 0.009510 | 0.012572 |
|  | Negative | 11.83049 | 3.895693 | 0.404728 | 0.065809 | 0.049793 |
| 1% | Positive | 0.000394 | 0.000741 | 0.003448 | 0.012650 | 0.015399 |
|  | Negative | 3.748166 | 1.306868 | 0.190816 | 0.049647 | 0.040670 |
| 5% | Positive | 0.002526 | 0.004296 | 0.010469 | 0.018774 | 0.020353 |
|  | Negative | 0.374346 | 0.182129 | 0.062877 | 0.033458 | 0.030762 |

Table 3 shows the skewness and kurtosis of the estimated coefficients, using (13) and (14), for the twenty sample specifications considered in Tables 1 and 2. The results in this table explain the pattern of findings in Tables 1 and 2. The estimated coefficient, viewed as a random variable under the null hypothesis, has strong negative skewness, inherited in turn from the negative skewness in the years-of-education variable. The negative skewness in the distribution of the years-of-education variable interacts with the asymmetry in the spread of SNP values (vastly more major allele observations than heterozygote or minor allele) to produce biases in the conventional p-values. For negative estimated betas, the conventional p-value understates Type I error, rejecting the null hypothesis when it should be accepted. For positive estimated betas, the conventional p-value sacrifices power by accepting the null hypothesis when it could be rejected with confidence.

**Table 3.** Skewness and kurtosis of GWAS regression coefficients under a mixture distribution: Years of education

| Sample Size: |  | 500 | 1000 | 5000 | 50000 | 100000 |
| --- | --- | --- | --- | --- | --- | --- |
| Minor allele frequency: |  |  |  |  |  |  |
| 0.1% | Skewness | -0.81933 | -0.57935 | -0.25909 | -0.08193 | -0.05794 |
|  | Kurtosis | 5.071641 | 4.035821 | 3.207164 | 3.020716 | 3.010358 |
| 0.5% | Skewness | -0.36198 | -0.25596 | -0.11447 | -0.03642 | -0.02575 |
|  | Kurtosis | 3.407713 | 3.203856 | 3.040771 | 3.004160 | 3.00208 |
| 1% | Skewness | -0.25201 | -0.1782 | -0.07969 | -0.02560 | -0.0181 |
|  | Kurtosis | 3.19976 | 3.09988 | 3.019976 | 3.002101 | 3.00105 |
| 5% | Skewness | -0.10559 | -0.07466 | -0.03383 | -0.01073 | -0.00759 |
|  | Kurtosis | 3.041966 | 3.020983 | 3.004343 | 3.000438 | 3.000219 |

## 5 Summary

This paper provides a new approach to estimating Bonferonni-corrected multiple-test p-values for regressions of individual genetic markers on a phenotypic variable. The current standard approach in computing coefficient p-values is to assume a normal distribution for the phenotypic variable, or to invoke the central limit theorem to justify the approximate normality of the coefficient estimate. Many phenotypic variables, particularly for social traits like income and education levels, have distributions which are far from normality. The central limit theorem, as we show, does not give reliable p-values for the types of sample sizes (and multiple-test numbers) used in genome-wide association studies (GWAS) with non-normally distributed phenotypic variables.

We suggest a new approach, based on fitting a Bernoulli-normal mixture distribution to the phenotypic variable (prewhitened with respect to any other confounding explanatory variables) before running univariate GWAS regressions. We derive the exact finite-sample distribution of GWAS regression coefficients p-values under this more flexible distributional assumption. We illustrate the magnitude of the p-value adjustment from our approach (relative to the conventional approach) with sample data on a commonly-used phenotypic variable with a notably non-normal distribution: years of education. The derived p-values differ markedly from the conventional, normality-based, p-values. This new approach allows for smaller samples sizes and lower minor allele frequencies since it does not rely on a large-sample central limit theorem approximation. We provide an implementation of the method in the R programming language, which is available as a standard R package from CRAN, along with user documentation.

## Acknowledgments

We would like to thank Philipp Koellinger, Robert Korajczyk, Richard Linner, Oliver Linton, Brian O’Kelly and Donal O’Neill for helpful comments. We wish to acknowledge support from the Science Foundation of Ireland.

## Appendix A: Technical Appendix

### 1 Introduction

This technical appendix discusses the computationally efficient procedures used in the R code, provides detailed derivations of the skewness and kurtosis of the estimated regression coefficient, and describes the years-of-education data set used in the paper.

### 2 The regression format

The p-value formula in GWRPV assumes that the regression coefficient 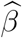 comes from a univariate genome-wide regression model, in which the influence of other confounding variates have been removed in a first-stage regression. The analyst begins with a raw phenotypic variable 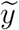 which potentially includes the influence of confounding variates *X*_*c*_, such as the dominant principal components of the genetic variants matrix. The raw phenotype 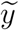 is first regressed on these confounding variates to remove their influence:

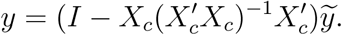

The prewhitened phenotypic variate *y* is then used throughout.

For a given SNP, the analyst has regressed the prewhitened phenotypic variable *y* on a single nucleotide polymorphism (SNP) *x*:

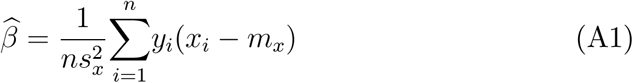

with sample {*y*_*i*_, *x*_*i*_}_*i*=1,*n*_, where *x*_*i*_ = 0 if the SNP is a major homozygote, *x*_*i*_ = 1 if the SNP is a heterozygote, *x*_*i*_ = 2 if the SNP is a minor homozygote. The inputs for computing the p-value of a given coefficient estimate are the five parameters of the mixture distribution, *µ*_*a*_, *σ*_*a*_, *µ*_*b*_, *σ*_*b*_, *p*, the number of each of the three values for the independent variable, *n*_0_, *n*_1_, *n*_2_, and the estimated coefficient, 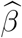.

The model assumes that *y* has a Bernoulli-normal mixture distribution. For notational simplicity, the main paper and this technical note also assume that *y* is standardized in the regression, so that *E*[*y*] = 0 and *E*[*y*^2^] = 1. However this assumption is only for notational simplicity; the analyst need not standardize *y* before running the regression (A1) or the GWRPV program. Under the assumed distribution, *E*[*y*] = *pµ*_*a*_ + (1 − *p*)*µ*_*b*_ and 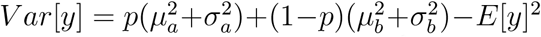. The GWRPV program takes the inputted regression coefficient 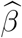 and, using the inputted parameters to compute *Var*[*y*], scales the inputted regression estimate by *Var*[*y*]^−1/2^. This has the effect of replacing the regression model with the equivalent one with standardized *y*. This linear transform of the regression model does not affect the p-value of the regression coefficient but simplifies the calculations.

The sample average sum of squares of *x* is:

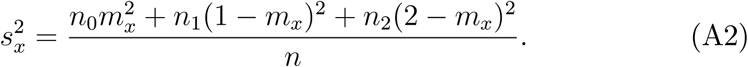

In running the regression, it is acceptable to subtract one from *x* and rescale so that *x* = −1, 0, 1 since this does not impact 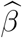 or 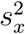. For notational simplicity we will assume *x* = 0, 1, 2.

### 3 Computing the exact p-value

Consider the case 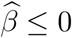. In order to determine the p-value, one considers this observed 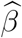 estimate and finds the cumulative probability of random outcomes which would give this value or less. (If 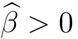 the p-value is computed as one minus this probability.) The cumulative distribution of an estimate 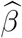 under the null hypothesis *β* = 0 is:

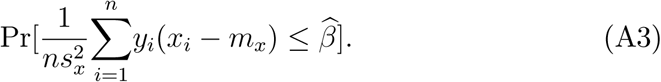

Each of the three integers *n*_0_, *n*_1_ and *n*_2_, decomposes into two (unobserved) integers: the number of realizations of the dependent variable *y*_*i*_ with the conditional normal distribution with mean and standard deviation {*µ*_*a*_, *σ*_*a*_} or {*µ*_*b*_, *σ*_*b*_}. We denote these integer realizations with a double subscript, {*n*_0*a*_, *n*_1*a*_, *n*_2*a*_}_*h*_,and {*n*_0*b*_, *n*_1*b*_, *n*_2*b*_}_*h*_, *h* = 1, *m* where the first subscript refers to the minor allele frequency, 0, 1, or 2, and the second subscript refers to the (unobserved) number of realizations of {*µ*_*a*_, *σ*_*a*_} or {*µ*_*b*_, *σ*_*b*_}, depending upon whether the Bernoulli random variable *λ* equals one or zero. Note that *m* = (*n*_0_ +1)(*n*_1_ +1)(*n*_2_ +1). Since the second set {*n*_0*b*_, *n*_1*b*_,*n*_2*b*_} consists of the remainders *n*_0*b*_ = *n*_0_ − *n*_0*a*_, *n*_1*b*_ = *n*_1_ − *n*_1*a*_ and *n*_2*b*_ = *n*_2_ − *n*_2*a*_, it is sufficient to index with {*n*_0*a*_, *n*_1*a*_, *n*_2*a*_}. Let *P*_*h*_, *h* = 1, *m* denote the probabilities of all these unobserved potential outcomes {*n*_0*a*_, *n*_1*a*_, *n*_2*a*_}_*h*_. The probability of the *n*–sum (A3) can be written as a probability-weighted *m*–sum over all the potential outcomes:

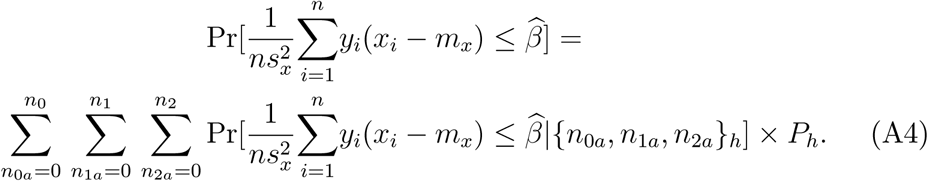

The probabilities of each potential outcome *P*_*h*_ = Pr[{*n*_0*a*_, *n*_1*a*_, *n*_2*a*_}_*h*_], can be found from the binomial formula (for three independent binomials).

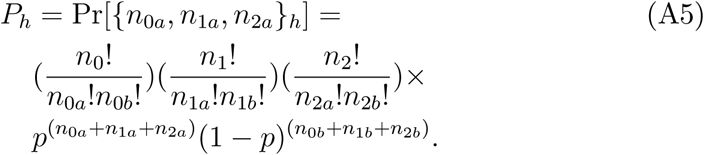

The formula (A5) consists of a potentially very large number, 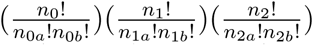 multiplied by a potentially extremely small number, 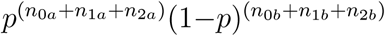. To prevent numeric overflow, the program computes the log of each of these two terms, sums the logs, and then takes the antilog.

### 4 Trimming the set of feasible combinations

The p-value formula (A4) requires a sum over the set of outcomes from three independent binomials with *n*_0_, *n*_1_ and *n*_2_ draws, giving a total of *m* = (*n*_0_ + 1)(*n*_1_ + 1)(*n*_2_ + 1) terms. If all three values *n*_0_, *n*_1_, *n*_2_ are very large (or even if only two are very large and the third is moderately large) this can be an extremely large number of terms to compute in the sum, and not computationally necessary. It is possible to shrink this computational burden substantially without sacrificing any estimation accuracy.

For each *h* = 1, *m*, rearrange (A4) listing the values associated with *n*_0_, *n*_1_, *n*_2_ separately:

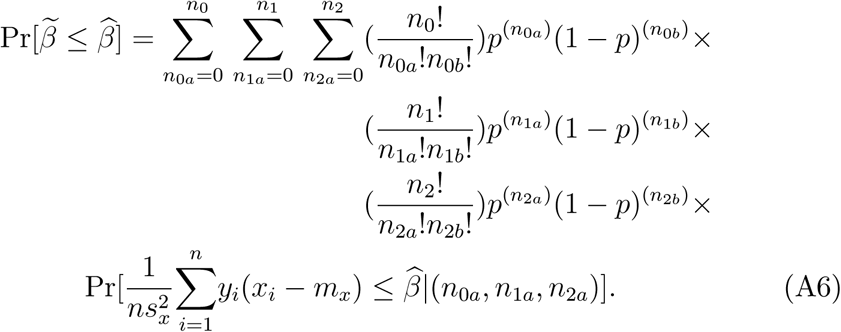

Each of the four multiplicative terms in (A6) is a probability and lies between zero and one, for every (*n*_0*a*_, *n*_1*a*_, *n*_2*a*_). Depending upon the values of *n*_0_, *n*_1_, *n*_2_, and *p*, the vast majority of the terms in (A6) are indistinguishable from zero.

To speed computation in the GWRPV programme, a range of beginning and/or ending terms in (A6) with extremely low cumulative probability are dropped from the sum. We divide the summation into two parts: a range of index values *A* which by construction covers virtually all of the total probability, and the complement set with total probability very close to zero. The separate computation interval for each of *n*_0*a*_, *n*_1*a*_, and *n*_2*a*_ is chosen using the univariate distributions. Let 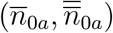 denote the upper and lower limits, with 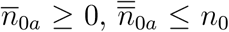 and 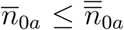. Let *δ* denote a small positive number (the GWRPV code uses *δ* = 10^−l6^ as the default value but this can be altered by the user). The range limits are chosen such that at most *δ* probability lies outside the range:

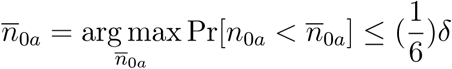

where of course Pr[*n*_0*a*_ < 0] = 0. Note that 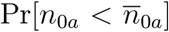 is the cumulative probability of a univariate binomial with parameter *p* (a trivial computation). Similarly for the upper limit on the range:

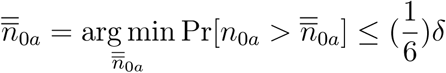

where of course Pr[*n*_0*a*_ > *n*_0_] = 0. The values for 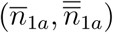 and 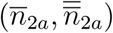 are chosen analogously. Since each of the ommitted tail ranges have probability less than or equal to 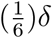, the set *A* has total probability greater than or equal to 1 − *δ*. The p-value is separated into the conditional probability given (*n*_0*a*_, *n*_1*a*_, *n*_2*a*_) ∈ *A* times the probability of (*n*_0*a*_, *n*_1*a*_, *n*_2*a*_) ∈ *A*, plus the remaining probability:

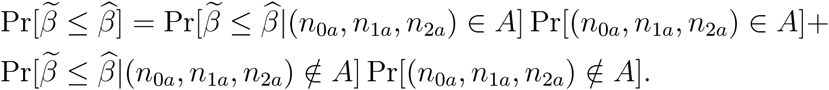

Using the restricted range A in (A6) in place of the full range:

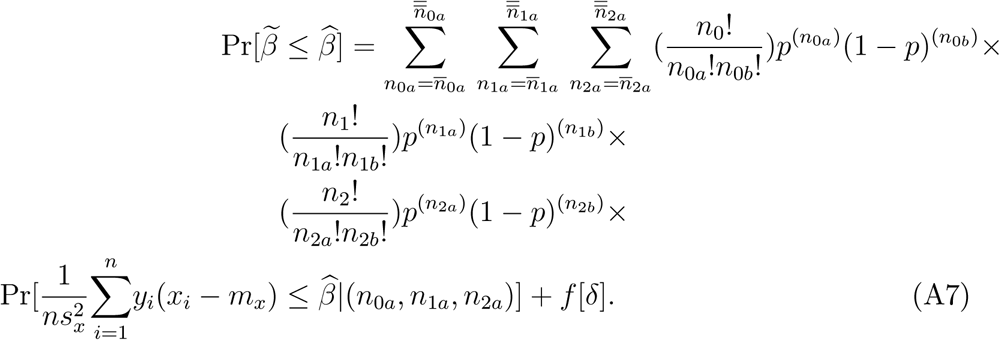

where *f*[*δ*] ≤ *δ*.

The trimming parameter is set by the parameter *logdelta* in the optional input file trimparameters.txt. *logdelta* is in log base 10 format and has a default value of − 16.

### 5 Using a normal approximation for the sample average phenotype of the major homozygote observations

If the sample size *n* is large and the minor allele frequency is low (so that the number of major allele observations *n*_0_ is large) then it may be possible to greatly speed computation by applying a central limit theorem approximation to the sample average of the phenotype values within the subset of major allele observations.

Recall that, for notational convenience, the observations are ordered by allele type. The beta coefficient 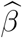 can be written as a linear combination of three random variables, the average of the phenotype over the major homozygote, heterozygote, and minor homozygote observations:

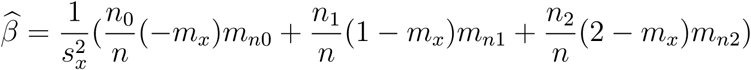

where

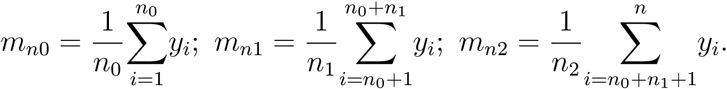

Since *y* has a Bernoulli-normal mixture distribution, if follows easily that *m*_*n*0_ is asymptotically normal for large *n*_0_. The GWRPV program computes the skewness and kurtosis of *m*_*n*0_ and measures how far they are from their normal-based values of zero and three. See Section 7 below for the formulas for *Skew*[*m*_*n*0_] and *Kurt*[*m*_*n*0_]. If (*Skew*[*m*_*n*0_])^2^ + (*Kurt*[*m*_*n*0_] – 3)^2^ < *nearnorm*, then the GWRPV program uses a normal approximation for *m*_*n*0_. This greatly shrinks the computation time of the program, since the computation loop (A6) need only run over *n*_1_ and *n*_2_, which tend to be much smaller than *n*_0_. The program has a default value of *nearnorm*= 10^−6^. The analyst can alter the value of *nearnorm* by inserting a different value of *lognearnorm* (in base 10 format) into trimparameters.txt.

### 6 Setting an upper limit on the computation sum

The GWRPV program has a built-in computation limit, *topsum*, designed to prevent an individual regression result from causing a long computation delay. Before running the main computation sum (A7) for each regression case 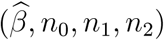, the program computes the number of terms in this computation sum, which is 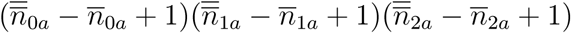. If this number is greater than *topsum*, and the skewness and kurtosis are not sufficiently close to normal values to invoke a normality-based p-value, then the program skips the p-value computation for that 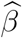 and procedes to the next regression case. A value of −999.9 is inserted in place of the p-value in the output file pvalues.txt to flag that the p-value computation has been skipped. The analyst can alter the value of *topsum* by inserting a different value of *logtopsum* (in base 10 format) into trimparameters.txt. The code has a default value of *topsum*= 10^8^.

### 7 The skewness and kurtosis of the estimated coefficient

In addition to computing the p-value, the GWRPV programme computes the skewness and kurtosis of the estimated regression coefficient, under the assumed mixture distribution of the dependent variable and under the assumed null hypothesis that the true coefficient equals zero. That is, the programme computes:

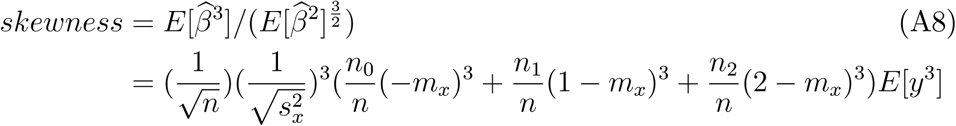

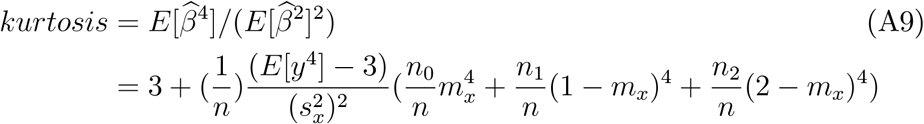

under the model’s assumptions. The first equalities in (A8) and (A9) are definitional; the second equalities will be derived here. Note that *skewness* → 0 and *kurtosis* → 3 as *n* → ∞.

Recall the definition of 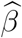, where we order the observations according to SNP allele type:

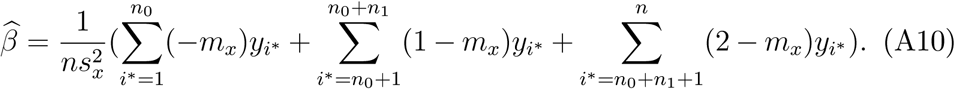

The three additive terms in (A10) will be denoted *a*, *b* and *c*. Since *y*_*i*_ is independently distributed across *i* and *E*[*y*_*i*_] = 0, we have *E*[*a*^*p*^*b*^*q*^*c*^*r*^] = *E*[*a*^*p*^]*E*[*b*^*q*^]*E*[*c*^*r*^] and *E*[*a*^*p*^*b*^*q*^*c*^*r*^] = 0 if *p*, *q* or *r* equals one. First consider the value in the numerator of both the skewness and kurtosis formulas (A8) and (A9). Writing the square using *a*, *b* and *c*:

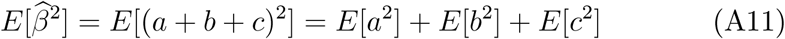

since all other terms in the product have at least one unit power of *a*, *b*, or *c* and so have zero expectation. Expanding out *E*[*a*^2^] and dropping terms which have a unit power of *y*_*i*_ for some *i*:

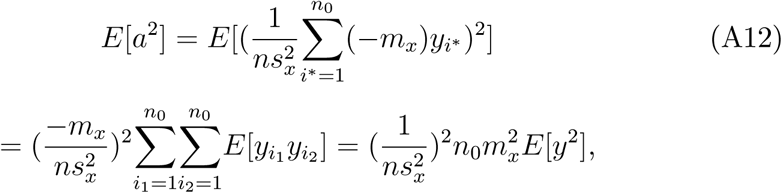

and recall that *E*[*y*^2^] = 1 since this variable is standardized. Repeating (A12) for *E*[*b*^2^] and *E*[*c*^2^]:

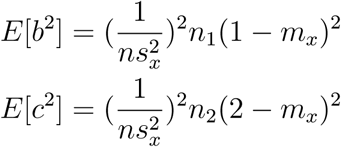

Inserting the expressions for *E*[*a*^2^], *E*[*b*^2^] and *E*[*c*^2^] into (A11):

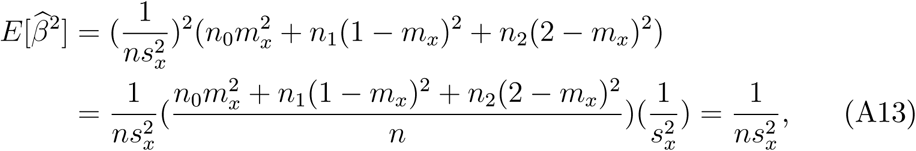

substituting the expression (A2) for 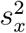 and cancelling from the numerator and denominator.

Next, we turn to the numerator in the definition of skewness (A8). Using (A10) gives:

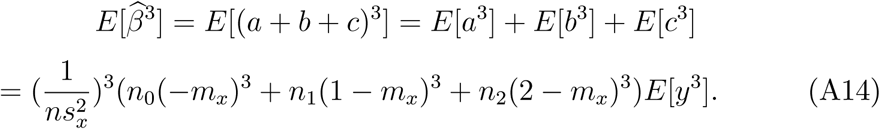

Taking the numerator divided by the denominator to get skewness:

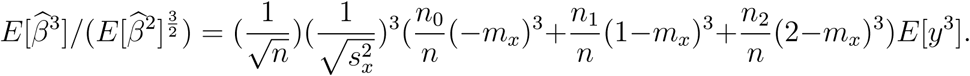

Following the same procedure as above, now for the computation of the numerator in the expression for kurtosis (A9):

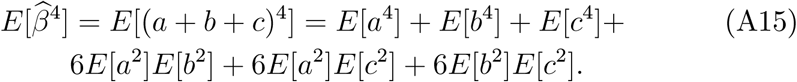

Consider first the term *E*[*a*^4^]. Expanding this out using the definition of a in (A10):

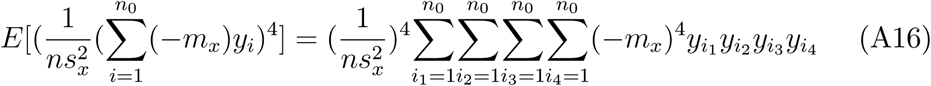

Dropping all terms in (A16) which include a unit power of *y*_*i*1_, *y*_*i*2_, *y*_*i*3_, or *y*_*i*4_ gives *n*_0_ terms with *i*_1_ = *i*_2_ = *i*_3_ = *i*_4_ and *n*_0_(*n*_0_−1) terms for each of the three cases {*i*_1_ = *i*_2_, *i*_3_ = *i*_4_, *i*_1_ ≠ *i*_3_}, {*i*_1_ = *i*_3_, *i*_2_ = *i*_4_, *i*_1_ ≠ *i*_2_}, {*i*_1_ = *i*_4_, *i*_2_ = *i*_3_, *i*_1_ ≠ *i*_2_}. The first set of terms all have the same individual term value of 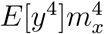 and similarly the other three sets of terms all have individual term values of 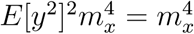. Summing all sets of terms:

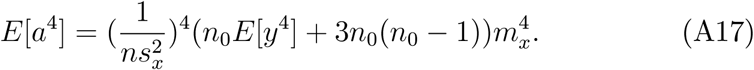

Repeating exactly the same steps to generate (A17) for *b* and *c*:

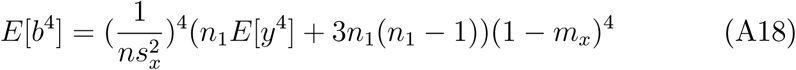

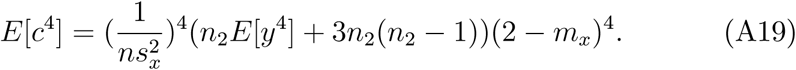

Inserting the expressions for *E*[*a*^2^], *E*[*b*^2^], *E*[*c*^2^], *E*[*a*^4^], *E*[*b*^4^], and *E*[*c*^4^] into (A15) gives:

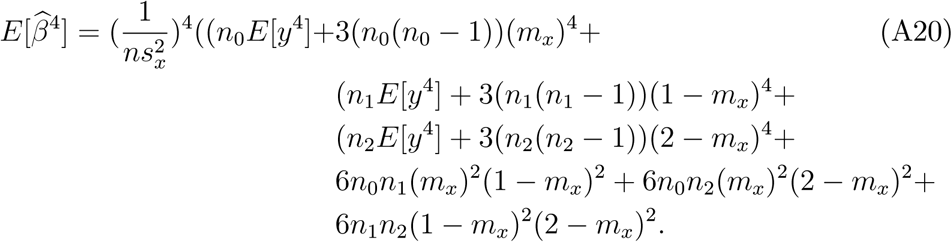

In order to simplify (A20) it is necessary to cancel out 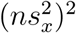 from a collection of terms in the the numerator. First, rewriting 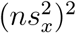 using the expression (A2) for 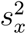:

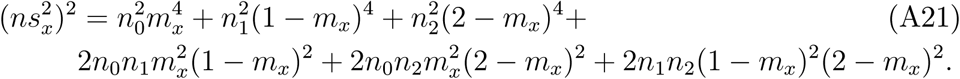

Gathering terms in (A20) that match those in (A21) inside the first curly bracket and the remainders (which all multiply (*E*[*y*^4^]−3)) inside the second curly bracket:

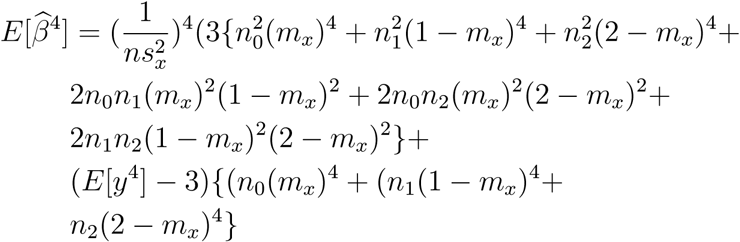

Dividing by 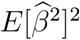 using (A13) and (A21) gives 3 plus a remainder:

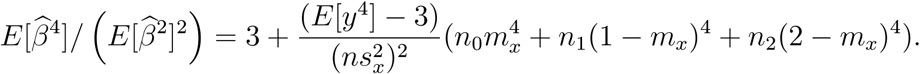

Next we derive the skewness and kurtosis of the Bernoulli-normal mixture distribution. We only consider the standardized case *E*[*y*] = 0 and *E*[*y*^2^] = 1. Recall that *y* = *λ*(*µ*_*a*_ +*σ*_*a*_*z*_*a*_)+(1−*λ*)(*µ*_*b*_ +*σ*_*b*_*z*_*b*_) where *λ* is a Bernoulli random variable with probability *p* that *λ* = 1 and *z*_*a*_, *z*_*b*_ are independent standard normal variates. Note that *λ*(1 − *λ*) = 0 and *λ*^2^ = *λ*^3^ = *λ*^4^ = *λ*. Taking the cube, dropping all terms which include *λ*(1 − *λ*), and simplifying powers of *λ* and (1 − *λ*):

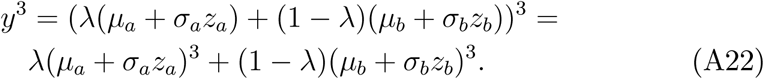

The third moment of a normal variate with mean *µ* and standard deviation *σ* is *µ*^3^ + 3*µσ*^2^. Taking the expectation of (A22) gives:

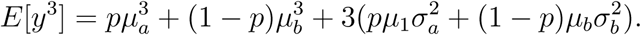

For the kurtosis of *y*, take the fourth power of *y*, drop all terms which include *λ*(1 − *λ*), and simplify powers of *λ* and (1 − *λ*):

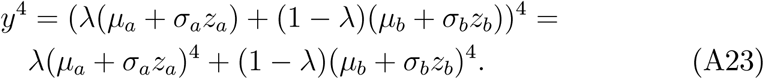

The fourth moment of a normal variate with mean *µ* and standard deviation *σ* is *µ*^4^ + 6*µ*^2^*σ*^2^ + 3*σ*^4^. Taking the expectation of (A23) gives:

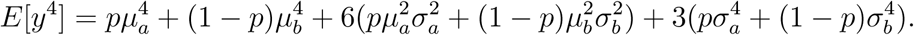

Finally, we derive the skewness and kurtosis of *m*_*n*0_, which is an average of *n*_0_ independent observations of *y*.Finding the second, third and fourth moments:

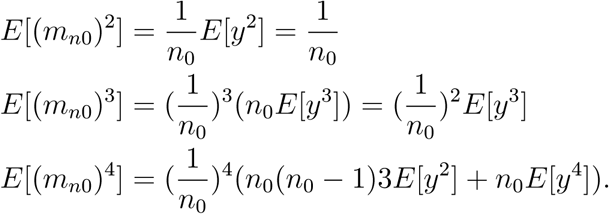

Taking ratios as in (A8) and (A9) to get skewness and kurtosis:

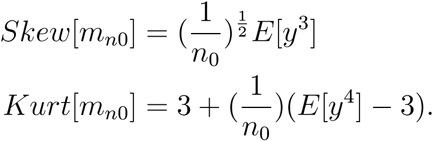

### 8 Years-of-education data description

The first two columns of Table A-1 below reproduce two rows from *Table 1: Educational Attainment of the Population 18 Years and Over, by Age, Sex, Race, and Hispanic Origin: 2015* in Current Population Survey Data on Educational Attainment (U.S. Census Bureau (2015)). We choose the subsample "U.S. white males ages 25 and greater" from that data source, which is row 25 of their *Table 1*. The [white/male/age 25 and over] sub-sample has 81,913 observations. Row three of Table A-1 below transforms the qualitative categories into a quantitative variable. There are a few minor subjective judgements in transforming the survey categories into quantitative years-of-education. The final column shows the frequency distribution of the data.

**Table A-1.**
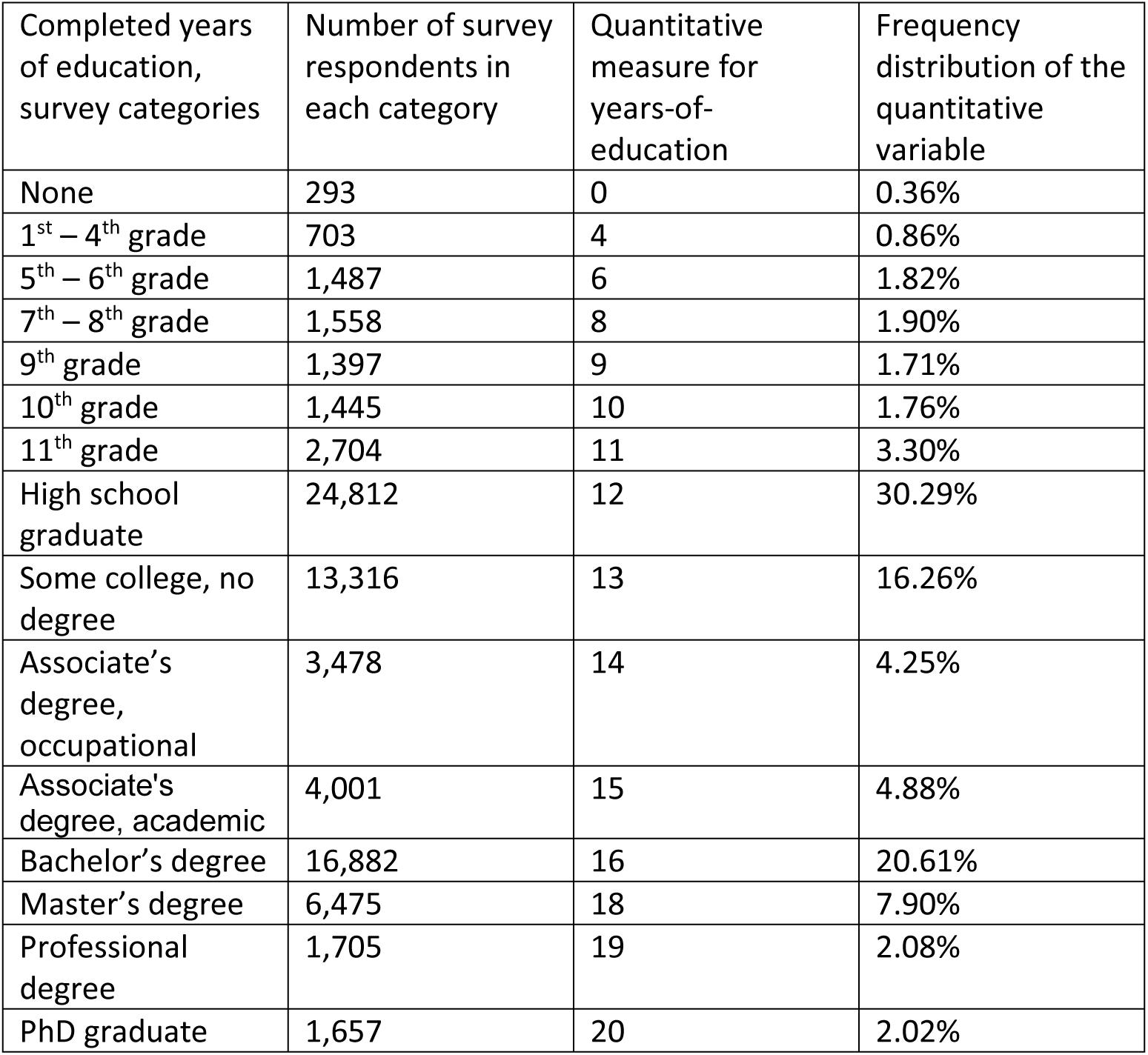
Years of education of U.S. white males, ages 25+

## Appendix B: Guide to the gwrpv program in the R package gwrpvr

### 1 Introduction

The Genome-Wide Regression P-Value (`gwrpv`) program computes the sample probability value (p-value) for the estimated coefficient from a standard genome-wide univariate regression. It computes the exact finite-sample p-value under the assumption that the measured phenotype (the dependent variable in the regression) has a known Bernoulli-normal mixture distribution. This appendix provides instructions for using the `gwrpv` program contained in the R package `gwrpvr` version 1.0.

### 2 Preliminary Research Steps

The `gwrpv` program only has added value if the phenotypic variable in the genome-wide regression study does not follow a normal distribution. Before running the `gwrpv` program, the analyst should compute the sample skewness and kurtosis of the phenotypic variable, and apply the Jarque-Bera test for normality. The Jarque-Beta test is provided in most statistical software packages; for example, *ajb.norm.test* in the normtest package in R (see ftp://cran.r-project.org/pub/R/web/packages/normtest/normtest.pdf) gives a standard implementation. If normality is not rejected by the Jarque-Bera test, the `gwrpv` routine is not appropriate; conventional normality-based p-values should be used instead.

In order to run the program, the analyst must fit a normal mixture distribution to the phenotypic variable. This can be done for example using the `normalmixEM` command in the mixtools library in R (see Benaglia et al. (2009)). The `gwrpv` program requires the six parameters of the fitted mixture distribution, *µ*_*a*_, *σ*_*a*_, *µ*_*b*_, *σ*_*b*_, *p*_*a*_ and *p*_*b*_. The parameters *µ*_*a*_, *σ*_*a*_ are the mean and standard deviation given that the Bernoulli random variable equals one, and *µ*_*b*_, *σ*_*b*_ are the mean and standard deviation given that the Bernoulli random variable equals zero. The parameter *p*_*a*_ is the probability that the Bernoulli random variable equals one and *p*_*b*_ is the probability that the Bernoulli random variable equals zero.

Before running the `gwrpv` program, the analyst must run a standard set of genome-wide regressions. That is, for each single-nucleotide polymorphism (SNP) in a large data set, the analyst has regressed a phenotypic variable *y*, observed over *n* individuals, on the realized SNP values across the individuals. The univariate regression has been run separately for each SNP in the data set, resulting in a very large number (up to tens of millions) of individual regression coefficients, one per SNP. This can be done using the plink package of routines (see Purcell, et al. (2007)). The `gwrpv` program can use as input the regression coefficient outputs of plink.

The `gwrpv` program allows the analyst to find p-values for any reasonable number of regression coefficients. The number of candidate regression coefficients should be a relatively small number, roughly in the range one to one thousand. It is not appropriate to compute adjusted p-values on the entire universe of tens of millions of regression coefficients, due to the processing-time demands of computing each exact finite-sample p-value. The candidate coefficients should be preselected by the analyst, based for example on the magnitude of their t-statistics, as the ones most likely to have statistically significant p-values. A sensible cutoff is to restrict the analysis to regression coefficients with a t-statistic greater than 3.5 in magnitude. For convenience we have also provided a batch version of the program (`gwrpv_batch`), which can process the results from a set of regressions.

### 3 Inputs and Outputs

The `gwrpv` program requires a number of inputs. The first required set of inputs are the six values (*µ*_*a*_, *σ*_*a*_, *µ*_*b*_, *σ*_*b*_, *p*_*a*_, *p*_*b*_). The inputs *µ*_*a*_ and *µ*_*b*_ can have any real values; *σ*_*a*_ and *σ*_*b*_ must be positive; *p*_*a*_ and *p*_*b*_ both must lie between zero and one, and must sum to one.

There are also a set of trimming parameters for efficient computation of the p-values (see Section 5 below), each of these is set to default values, which can be overridden.

The second required set of inputs are four values^1^. The first is the set of candidate regression coefficient estimates for the regressions being analyzed. The second, third, and fourth values must give the number of major heterozygote, homozygote, and minor heterozygote observations from each of the corresponding regression samples. The three number-of-observations columns must all be non-negative integers; the beta estimates can have any real values.

The `gwrpv` program outputs the p-value of the regression coefficient based on the mixture distribution and some supplementary statistics, including the skewness and kurtosis of the coefficient estimate, based on the assumed mixture distribution for the dependent variable.

### 4 Procedure for Running the Program

This short section lists the steps needed to run the program. The `gwrpv` program is contained in the R package `gwrpvr`, which is available as standard from CRAN. The current version of the program is v1.0.

In your favourite R environment install the package `gwrpvr`.

~~~
> install.packages("gwrpvr")
> library(gwrpvr)
~~~

To see the associated help file for the `gwrpv` program with the required parameters run

~~~
> help("gwrpv")
~~~

The following is an example of how to use `gwrpv`, having initially calculated the inputs as per Sections 2 and 3.

~~~
> beta <- 6.05879 # candidate regression cooefficient estimate
> n0 <- 499 # number of major heterozygote observations
> n1 <- 1 # number of major homozygote observations
> n2 <- 0 # number of minor heterozygote observations
> mua <- 13.87226 # mean of the mixture distribution given that the Bernoulli random variable equals zero
> siga <- 2.58807 # stdev of the mixture distribution given that the Bernoulli random variable equals zero
> mub <- 4.62829 # mean of the mixture distribution given that the Bernoulli random variable equals one
> sigb <- 2.51803 # stdev of the mixture distribution given that the Bernoulli random variable equals one
> pa <- 0.96544 # pa is the probability that the Bernoulli random variable equals one
> pb <- 0.03456 # 1 - pa
> g <- gwrpv(beta, n0, n1, n2, mua, siga, mub, sigb, pa, pb)
> g$pvalue # display the p-value
> g # display all the output statistics
~~~

#### 4.1 The skewness and kurtosis of the estimated coefficient

In addition to computing the exact p-value, the `gwrpv` programme computes the skewness and kurtosis of the estimated regression coefficient, under the assumed mixture distribution of the dependent variable, and assuming the null hypothesis that the true coefficient equals zero.

Following on from the sample input parameters in the earlier example we can retrieve the output skewness and kurtosis as follows:

~~~
> g <- gwrpv(beta, n0, n1, n2, mua, siga, mub, sigb, pa, pb)
> g$pvalue # display the p-value
> g$skew # display the skewness
> g$kurt # display the kurtosis
~~~

### 5 Efficient Computation Procedures

If applied naively, the p-value computation in the model requires a sum over a potentially very large number of terms. The `gwrpv` programme uses efficient computation procedures to minimize run-time, while maintaining a high degree of accuracy in the p-value computation. There are three parameter inputs which control these computation features in the programme: *logdelta*, *lognearnorm* and *logtopsum*. All three parameters are real numbers inputted in log base ten format.

The parameter *logdelta* controls the trimming of the three univariate binomial distributions, each binomial distribution corresponding to the sample of one allele type (see the Technical Appendix for analytical details). *logdelta* has a default value of -16, which means that the p-value computation is only accurate for 16 decimal places. The analyst can increase or decrease *delta* by changing the optional parameter value; a larger-magnitude negative delta will result in a more accurate computation and slower run-time. *logdelta* must be a negative number, in log base ten format, so that 10^*logdelta*^ is less than one.

The parameter *lognearnorm* is used to determine whether to use a triple computation loop over *n*0, *n*1 and *n*2 observations, or apply the central limit theorem to approximate the distribution of the sample average phenotype of the major homozygote observations. This approximation eliminates the need to loop over *n*0, giving a double loop over *n*1 and *n*2 only. The program computes the skewness and kurtosis of the sample average phenotype of the major homozygote observations. If the sum of squared difference of skewness and kurtosis from their normal distribution values is less than nearnorm, then the `gwrpv` program uses the central limit theorem to eliminate the major homozygote observations from the computation loop.

The final parameter, *logtopsum*, ensures that the `gwrpv` programme does not spend too long computing a p-value. Before running the main computation sum for each regression case 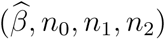, the program computes the number of terms in this computation sum. If this number is greater than *logtopsum*, the program skips the p-value computation for that 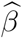 and proceeds to the next regression case, and a value of −999.9 is inserted in place of the p-value in the output to indicate that the computation has been skipped.

#### 5.1 Inputting the control parameters for efficient computation

In the `gwrpv` program we provide parameters to faciliate efficient computation. These are the control parameters described above, the three real numbers, *logdelta*, *lognearnorm* and *logtopsum*. All three of these inputs must be provided in log base ten, so that (−16, −5, 8) means that *δ* is set at 10^*−*16^ and *topsum* is 10^8^.

By default we set values for these three parameters (i.e., logdelta=-16, lognearnorm=-5, and logtopsum=8). These do not need to be explicitly passed in unless you want to override them. For example, the following will result in the same output as the earlier example. Here we are explicitly setting the trimming parameters..

~~~
> g <- gwrpv(beta, n0, n1, n2, mua, siga, mub, sigb, pa, pb, logdelta=-12, lognearnorm=-5, logtopsum=8)
> g$pvalue
~~~

### 6 Batch mode

If one wishes to compute p-values for mutiple regressions there is a batch version of the function, `gwrpv_batch`. The following are examples of its use. They illustrate how the results of each regression are presented to `gwrpv_batch` as a list of lists.

~~~
# create a list of the beta’s
> beta <- c(6.05879, -6.05879, 2.72055, -2.72055, 1.93347, -1.93347, 0.88288, -0.88288, 4.28421, -4.28421)
# create a list of the number of major heterozygote observations
> n0 <- c(499, 499, 495, 495, 490, 490, 451, 451, 998, 998)
# create a list of the number of major homozygote observations
> n1 <- c(1, 1, 5, 5, 10, 10, 48, 48, 2, 2)
# create a list of the number of minor heterozygote observations
> n2 <- c(0, 0, 0, 0, 0, 0, 1, 1, 0, 0)
# create the list of lists
> myregresults <- list(beta = beta, n0 = n0, n1 = n1, n2 = n2)
> g <- gwrpv_batch(myregresults,13.87226,2.58807,4.62829,2.51803,0.96544,0.03456)
~~~

In the second example we illustrate how to load the regression results from a comma separated file. Connor & O’Neill (2017) describe an illustrative sample data set of regressions to which the Genome-Wide Regression P-Value method is applied. The R package `gwrpvr` contains a folder called `data`/ in which this data set is provided. The data is contained in the file named `regresults.csv`

~~~
# alternatively the regression results may be contained in a .csv file # let’s call the file "regresults.csv"
# assuming four comma separated columns with a single header line
# containing columns names: beta,n0,n1,n2
# readr is a handy package to read in csv files, install this if not in your environment
> install.packages(’readr’)
# load the readr package into your environment
> library(readr)
# use the read_csv function from the readr package to load in the csv
> myregresults <- read_csv("data/regresults.csv")
> g <- gwrpv_batch(myregresults,13.87226,2.58807,4.62829,2.51803,0.96544,0.03456)
~~~

## Footnotes

1 There is a batch version of the function gwrpv batch, to which a list of candidate regression coefficients whose adjusted p-values are to be computed are passed.

